# PDEδ inhibition impedes the proliferation and survival of human colorectal cancer cell lines harboring oncogenic KRas

**DOI:** 10.1101/377259

**Authors:** Christian H. Klein, Dina C. Truxius, Holger A. Vogel, Jana Harizanova, Sandip Murarka, Pablo Martín-Gago, Philippe I. H. Bastiaens

**Author notes:** Correspondence to: Mail, Telefon.

## Abstract

**Novelty and Impact:** The ‘undruggable’ KRas is a prevalent oncogene in CRC with poor prognosis. In hPDAC cells pharmacological targeting of PDEδ affects oncogenic KRas signaling, but it remained unclear whether this approach is transferable to other cancer cells. Here, we show that genetic and pharmacologic PDEδ inhibition also impedes the proliferation of oncogenic, but not wild-type KRas bearing CRC cells indicating that PDEδ inhibition is a specific tool for targeting growth of oncogenic KRas bearing CRC.

**Abstract:** Ras proteins, most notably KRas, are prevalent oncogenes in human cancer. Plasma membrane localization and thereby signaling of KRas is regulated by the prenyl-binding protein PDEδ. Recently, we have reported the specific anti-proliferative effects of PDEδ inhibition in KRas-dependent human pancreatic ductal adenocarcinoma cell lines. Here, we investigated the proliferative dependence on the solubilizing activity of PDEδ of human colorectal cancer (CRC) cell lines with or without oncogenic KRas mutations. Our results show that genetic and pharmacologic interference with PDEδ specifically inhibits proliferation and survival of CRC cell lines harboring oncogenic KRas mutations whereas isogenic cell lines in which the KRas oncogene has been removed, or cell lines with oncogenic BRaf mutations or EGFR overexpression are not dependent on PDEδ. Pharmacological PDEδ inhibition is therefore a possible new avenue to target oncogenic KRas bearing CRC.

## Introduction

Ras proteins, most prevalent isoform KRas4B [1], are mutated in around 30 % of all human cancers [2] and especially frequent in pancreatic, colorectal and lung tumors [3]. Oncogenic mutations retain Ras in a constitutively active conformation [4], causing sustained activation of downstream signaling cascades leading to increased proliferation and survival [5]. Signal transduction from active KRas is dependent on its plasma membrane (PM) localization [6]. Despite a polybasic stretch and a farnesyl motif at the C-terminus of KRas conferring association to the negatively charged inner leaflet of the PM, this localization is compromised by endocytosis and entropy-driven re-equilibration to all endomembranes. The guanine nucleotide dissociation inhibitor (GDI-) like solubilization factor – PDEδ – counters this re-equilibration by binding the farnesyl-tail of KRas, thereby effectively increasing diffusion in the cytosol. KRas is then released in the perinuclear area by activity of the small GTPase Arl2 [7, 8] and electrostatically trapped and enriched on the recycling endosome (RE). This concentrated KRas on the RE is transported back to the PM via vesicular transport to maintain its enrichment there [8]. Interference with the solubilizing PDEδ functionality stalls this spatial cycle that maintains KRas concentration on the PM [8], thereby impairing KRas signaling [8, 9]. These findings led to the development of various small-molecule inhibitors of PDEδ based on different chemical scaffolds (Deltarasin, Deltazinone 1, Deltasonamide 1 & 2) that all competitively interact with the farnesyl-binding pocket [10-12]. In previous studies, we investigated the applicability of theses inhibitors on human pancreatic cancer cell lines since the majority (90 %) of pancreatic tumors harbor oncogenic KRas mutations [3, 13]. All three inhibitor classes reduced cell proliferation of KRas-dependent human pancreatic ductal adenocarcinoma cells (hPDACs), whereas KRas-independent or wild-type KRas harboring hPDACs were less affected [10-12].

Here, we expand the applicability of pharmacological PDEδ interference to colorectal cancer (CRC), another tumor class with prevalent (45 %) oncogenic KRas mutations [3]. To date, targeted therapy with monoclonal antibodies against EGFR, such as Cetuximab, is a major alternative to systemic cytotoxic chemotherapy in CRC [14]. However, therapy based on EGFR inhibition fails if oncogenic KRas [15, 16] or BRaf [16, 17] are expressed in CRC. To assess if PDEδ inhibition could be a possible new avenue to affect oncogenic KRas bearing CRC, we studied the dependence of CRC cell proliferation and survival on PDEδ activity. For this, we compared the effects of doxycycline-induced shRNA mediated down regulation of PDEδ to the effects of pharmacological interference with PDEδ activity in a panel of human CRC cell lines harboring distinct oncogenic mutations. We find a high correlation between the effects of pharmacological inhibition and shRNA-mediated PDEδ knock down on CRC proliferation and survival, where oncogenic KRas bearing CRC cells are highly compromised in cell proliferation and survival, whereas CRC cell lines in which the KRas oncogene was removed, or that harbor other oncogenic mutations, are hardly or not affected by PDEδ interference. Our findings suggest that PDEδ could be a valid therapeutic target for oncogenic KRas-driven colorectal cancer.

## Materials and Methods

### Cell culture

HCT-116 (ATCC American Type Culture Collection, Manassas, VA, USA), Hke3 (kind gift from Dr. Owen Sansom), Hkh2 (kind gift from Prof. Dr. Walter Kolch), DiFi (kind gift from Dr. Clara Montagut) and SW480 (ATCC) cell lines were maintained in DMEM (Dulbecco’s modified Eagle medium, Sigma-Aldrich Biochemie GmbH, Taufkirchen, Germany) supplemented with 10 % FCS (fetal calf serum; Pan-Biotech GmbH. Aidenbach, Germany), 2 mM L-glutamine (Sigma-Aldrich Biochemie GmbH) and 1 % NEAA (non-essential amino acids) (Sigma-Aldrich Biochemie GmbH), at 37°C and 5 % CO_2_ in a humidified incubator.

HT29 cells (ATCC) were maintained in Ham’s medium (Sigma-Aldrich Biochemie GmbH), supplemented with 10 % FCS and 1 mM L-glutamine (Sigma-Aldrich Biochemie GmbH), at 37°C and 5 % CO_2_ in a humidified incubator.

Cell line identity was validated by STR-profiling (DSMZ, Braunschweig, Germany) and all cell lines were routinely tested for mycoplasma.

### Small-molecule inhibitors

Deltarasin (Lot. No. 1) was purchased from Chemietek, Indianapolis, In, USA. Deltasonamide 2 was synthesized in-house as described previously [12].

### Virus production and generation of stable cell lines

Lentiviruses were produced and harvested as described previously, utilizing the most effective shRNA sequence against PDE6D (pLKO-PDE6D-572, see below) from a previous screen [10]. Viral supernatant, containing 10 μg/ml polybrene, was immediately used to infect target cells in 6-well plates at 50% confluence. After 24 h, lentivirus-containing supernatant was removed and fresh medium supplied, containing the appropriate amount of puromycin for selection. Puromycin tolerance was tested for all target cell lines prior to shRNA transduction.

*pLKO-shRNA-PDE6D-572:*

*sense:* 5’-CCGGGCACATCCAGAGTGAGACTTTCTCGAGAAAGTCTCACTCTGG AT GT GCTTTTT G-3’,

*antisense:* 5’-AATT CAAAAAGCACAT CCAGAGT GAGACTTT CTCGAGAAAGT CT CA CTCTGGATGTGC-3’

### Western Blot analysis

For PDEδ protein level analysis, whole cell lysates (WCL) were prepared after 72 h of doxycycline (200 ng/ml) induction as described previously [11]. For enrichment of Ras-GTP, 3xRaf-RBD pull down was executed. Recombinant GST-3xRafRBD [9] was expressed in E. Coli BL21DE3 by induction with 0.1 mM IPTG for 5 h after the culture reached an OD_600_ of 0.8. Afterwards, bacteria were harvested and lysed with bacterial lysis buffer (50 nM Tris-HCl, pH 7.5, 400 mM NaCl, 1mM DTT, 1 % Triton X-100, 1mM EDTA, supplemented with Complete Mini EDTA-free protease inhibitor (Sigma-Aldrich Biochemie GmbH) and bacterial lysates were stored at – 20°C. For the pull down, 700 μg crude bacterial lysate was incubated with magnetic GSH sepharose 4B beats for 2h at 4 °C on a rotating wheel and afterwards beats were re-equilibrated in cell lysis buffer (50 mM Tris-HCl pH 7.5, 200 mM NaCl, 10 % Glycerol, 2.5 mM MgCl2, 1 % Triton X-100, supplemented with Complete Mini EDTA-free protease inhibitor). Whole cell lysates were prepared after over night starvation in cell lysis buffer. 25 μg of WCL were used as “input control” to determine panRas, PDE6D and Cyclophilin B level, whereas 400 μg of WCL was subjected to GST-3xRaf-RBD, bound onto GSH sepharose 4B (GE) beats, pull down. After incubation for 30 min at 4 °C on a rotating wheel, beats were washed three times with cell lysis buffer. Then, bound Ras-GTP was eluted with SDS sample buffer for 10 min at 95 °C. Afterwards, SDS-polyacrylamide gel electrophoresis was carried out. Gels were blotted onto PVDF membrane (Immobilon, Millipore) and blocked for 1 h at room temperature with blocking buffer (LI-COR, Lincoln, NE, USA). The following antibodies were used for western blotting in the stated dilution: anti-PDE6D (Santa Cruz: sc-50260, 1:200), anti-Cyclophilin-B (Abcam: Ab16045, 1:3,000), anti-panRas (Calbiochem: OP40, 1:1,000) and matching secondary infrared antibodies IRDye 680 donkey anti rabbit IgG, IRDye 800 donkey anti mouse/goat IgG, (LI-COR, 1:10,000). Blots were scanned on a LI-COR Odyssey imaging system. Western blots were quantified using the Gel profiler plugin of ImageJ. Uncropped blots are shown in Supplementary figures 1 and 2.

### Clonogenic assays

Sparsely seeded cells (1–2 10^3^ per well) were maintained in a 6-well plate in the presence or absence of doxycycline (200 ng/ml). Doxycycline was applied 24 h after seeding. After ten days, cells were fixed and stained with 0.05 % (v/v) crystal violet (Sigma-Aldrich Biochemie GmbH) to visualize individual colonies. The quantification was performed using the analyze particle plug-in of ImageJ to extract total cell number and average colony size after utilizing a cell profiler pipeline to separate overlapping colonies.

### Real-time cell analyzer (RTCA)

RTCA measurements were performed using 16-well E-plates on a Dual Plate xCELLigence instrument (Roche Applied Science) in a humidified incubator at 37°C with 5 % CO2. The system measures the impedance-based cell index (CI), a dimensionless parameter which evaluates the ionic environment at the electrode/solution interface and integrates this information on the cell number [18]. Continuous impedance measurements were monitored every 15 min for up to 300 hours. Blank measurements were performed with growth medium. Depending on the cell line, 1 · 10^4^ - 2 · 10^4^ cells were plated in each well of the 16-well plates for short-term measurements and 0.75 - 2 · 10^3^ cells/well for long-term measurements. After seeding, cells were allowed to reach steady growth for 24 h before small-molecule inhibitor administration, whereas in case of cells stably expressing the inducible shRNA against PDEδ, doxycycline was directly applied to the wells of interest. In case of dose-dependent inhibitor measurements, the amount of DMSO was kept constant between the individual conditions and did not exceed 0.24 %. Cell indices were normalized to the time point of drug administration. For shRNA experiments no normalization was applied.

### Apoptosis assay

Apoptosis assays were performed on a LSR II flow cytometer (BD Bioscience, Heidelberg, Germany). For this, cells were seeded in 6-well plates at 2 · 10^5^ cells per well. Cells were treated with different concentrations of small-molecule inhibitors (Deltarasin or Deltasonamide 2) for 24 h. DMSO was used as a vehicle control. Subsequently, the supernatant was collected in FACS vials and the cells were washed with 1 mL PBS. Afterwards, cells were detached with 0.5 mL Accutase™ (EMD Millipore Corporation). The detached cells were re-suspended in 1 mL PBS and transferred to the respective FACS vials and centrifuged at 200 g for 5 min. The supernatant was discarded and the cells were washed twice with PBS. Cell pellets were re-suspended in 100 μl PBS containing 5 μl of 7-AAD (BD Bioscience). Samples were vortexed and incubated in the dark at RT for 15 min. Afterwards, 200 μL PBS were added and the samples transferred to fresh FACS vials through filter lids. The samples were measured within one hour after transfer using 488 nm as excitation wave length and the emission filter 695/40. Measurements were acquired and gated with the BD FACSDiva™ software.

## Results and Discussion

We studied the effects of genetic and pharmacological PDEδ interference in a cell panel containing six human CRC cell lines lacking or bearing distinct oncogenic mutations (table 1). While the SW480 cell line is homozygote for oncogenic KRas [19], HCT-116 cells contain one mutant and one wild-type KRas allele [20]. With the goal to create isogenic cell lines to HCT-116 that do not harbor oncogenic KRas, the Hke3 and Hkh2 cell lines were derived from HCT-116 by exchanging the mutant KRas allele with a non-transcribed KRas mutant (G12C) allele using homologous recombination [20]. However, the recombination was only successful in Hkh2, while Hke3 cells still contain an allele encoding oncogenic KRas that is expressed at lower levels (dosage effect mutant) [21]. In addition, we studied two CRC cell lines expressing wild-type KRas that have other oncogenic mutations. The HT29 cell line bears an oncogenic BRaf mutation (V600E) [17], an effector of Ras [22], whereas DiFi cells harbor an amplification of the EGFR gene accompanied with increased level of EGFR protein expression [23, 24] and are one of the few available cell models that are sensitive to anti-EGFR mAb treatment [25]. To study effects of PDEδ knock down on proliferation and viability, CRC cells were transduced with a lentivirus encoding a previously reported doxycycline-inducible short hairpin RNA (shRNA) sequence against PDEδ that is stably incorporated into their genome [10, 11]. shRNA expression was induced by doxycycline over several days and PDEδ protein levels were determined by western blot analysis at different time points in Hke3 cells. PDEδ levels decreased over time after doxycycline induction and a good knock down efficiency of >80 % was reached after 72 h (supplementary figure 1 A), which is consistent with the low protein turnover of PDEδ [9]. To now compare the amount of PDEδ expression levels of the different cell lines as well as to evaluate the knock down efficiency, western blot analysis of PDEδ protein levels was performed with/without doxycycline induction for 72 h (figure 1 A). All transduced cell lines showed a clear reduction in PDEδ protein levels upon doxycycline induction with respect to the corresponding control. Comparison of PDEδ expression levels in the non-induced CRC cell lines revealed that SW480 cells (homozygote for KRasG12V) exhibited the highest PDEδ level, whereas the KRas wild-type expressing HT29 cell line contained the lowest amount of PDEδ protein. Since PDEδ is necessary to maintain the PM localization of KRas and thereby its signaling activity [8, 9], we next investigated if there was a correlation between PDEδ expression and KRas activity among the different CRC cell lines. For this, we quantified the expression levels of PDEδ and Ras within the parental cell lines by western blot analysis (figure 1 B). The amount of GTP-loaded Ras was also quantified by specific precipitation of Ras-GTP from whole cell lysates using 3xRBD (three repeats of Ras binding domain of cRaf [9]) from cells that were serum-starved 24 hours prior to cell lysis. A strong correlation between PDEδ levels and total Ras expression (Pearson’s correlation coefficient of r^2^ = 0.974) as well as Ras activity (r^2^ = 0.949) became apparent, suggesting a dependence of oncogenic Ras activity on PDEδ expression levels.

**Figure 1.**
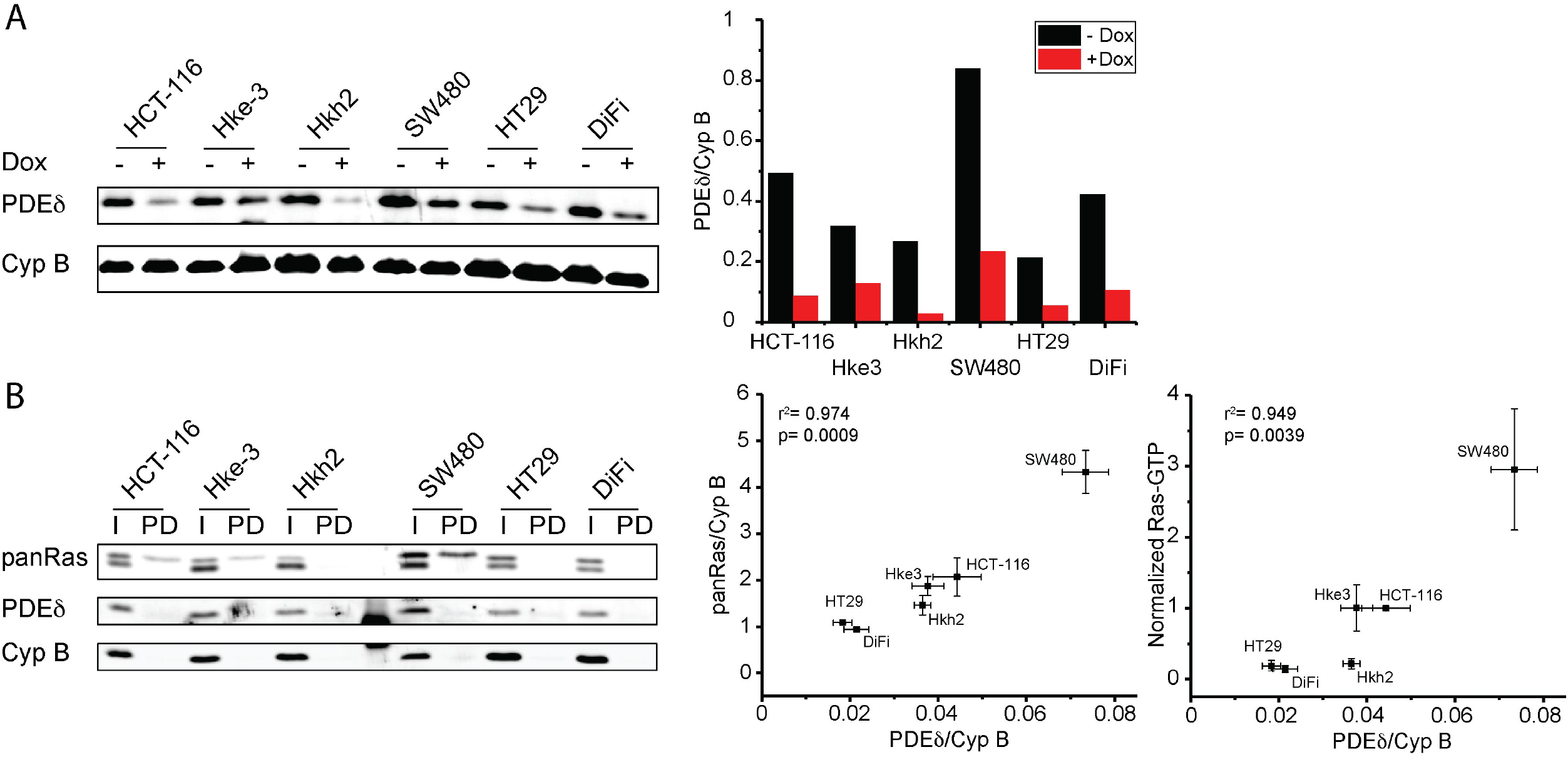
PDEδ and Ras levels in colorectal cancer cell lines. **(A)** Left: PDEδ protein level of distinct colorectal cancer cell lines in absence or presence of PDEδ shRNA induced by doxycycline after 72 h determined by western blot analysis. Cyclophilin B was used as loading control. Right bar graph: quantification of endogenous PDEδ levels of each cell line with (red) and without (black) doxycycline induction. **(B)** Left: PDEδ and panRas protein level (I) and Ras-GTP level (PD) of distinct CRC cell lines determined by western blot analysis. Cells were serum-starved 24 h before lysis and active Ras was enriched by 3xRaf-RBD pull-down. Middle and right: Correlation plots of PDEδ and panRas expression ± s.e.m of four biological replicates as well as PDEδ and active Ras levels ± s.e.m of four biological replicates (normalized to HCT-116 data). Pearson’s correlation analysis shows a high correlation of 0.974 and 0.949 between the respective expression levels.

**Table 1.** Overview of colorectal cancer cell lines used in this study including KRas mutation status as well as other relevant oncogenic mutations.

| Cell line | KRas status | Other onc. mutations |
| --- | --- | --- |
| SW480 | G12V//G12V | - |
| HCT-116 | G13D//wt | - |
| Hke3 | G13D//wt | - |
| Hkh2 | -//wt | - |
| HT29 | wt//wt | BRaf (V600E) |
| DiFi | wt//wt | EGFR overexpression |

We next performed clonogenic assays [26] to study the effect of doxycycline induced PDEδ knock down on proliferation and viability of the different CRC cell lines (figure 2 A). For this, CRC cells stably transduced with doxycycline-inducible shRNA against PDEδ were grown in the presence of doxycycline and the colony number and size was compared to that of untreated control after a growth period of ten days. Here, the number of colonies that remain after PDEδ knock down is a measure of cell viability, whereas the colony size is a measure of cell proliferation. Quantification of these two parameters (figure 2 B) showed significant growth-inhibition and viability reduction as a result of PDEδ knock down only in the oncogenic KRas bearing SW480, HCT-116 and Hke3 cell lines, but not in the HT29 cell line (oncogenic BRafV600E) and DiFi cells with EGFR overexpression, while the isogenic oncogenic KRas-lacking Hkh2 cell line showed only a minimal decrease in cell proliferation. A clear correlation between oncogenic KRas expression and viability as well as proliferation could be observed upon doxycycline induced PDEδ knock down in the CRC cells (figure 2 E, Pearson’s correlation coefficient of r^2^ = 0.909). A clear separation of CRC cells with and without KRas mutation became also apparent. Where SW480 cells, in which both KRas alleles are mutated, exhibited the strongest reduction in proliferation and cell viability. Both HCT-116 and Hke3 cells (heterozygote KRasG13D) showed a comparable reduction in cell proliferation upon PDEδ knock down, whereas the viability of the low oncogenic KRas-expressing Hke3 was substantially less affected. In contrast, the wild-type KRas bearing CRC cells (HT29, Hkh2) were hardly affected in their viability and proliferation and DiFi cells were not affected at all.

**Figure 2.**
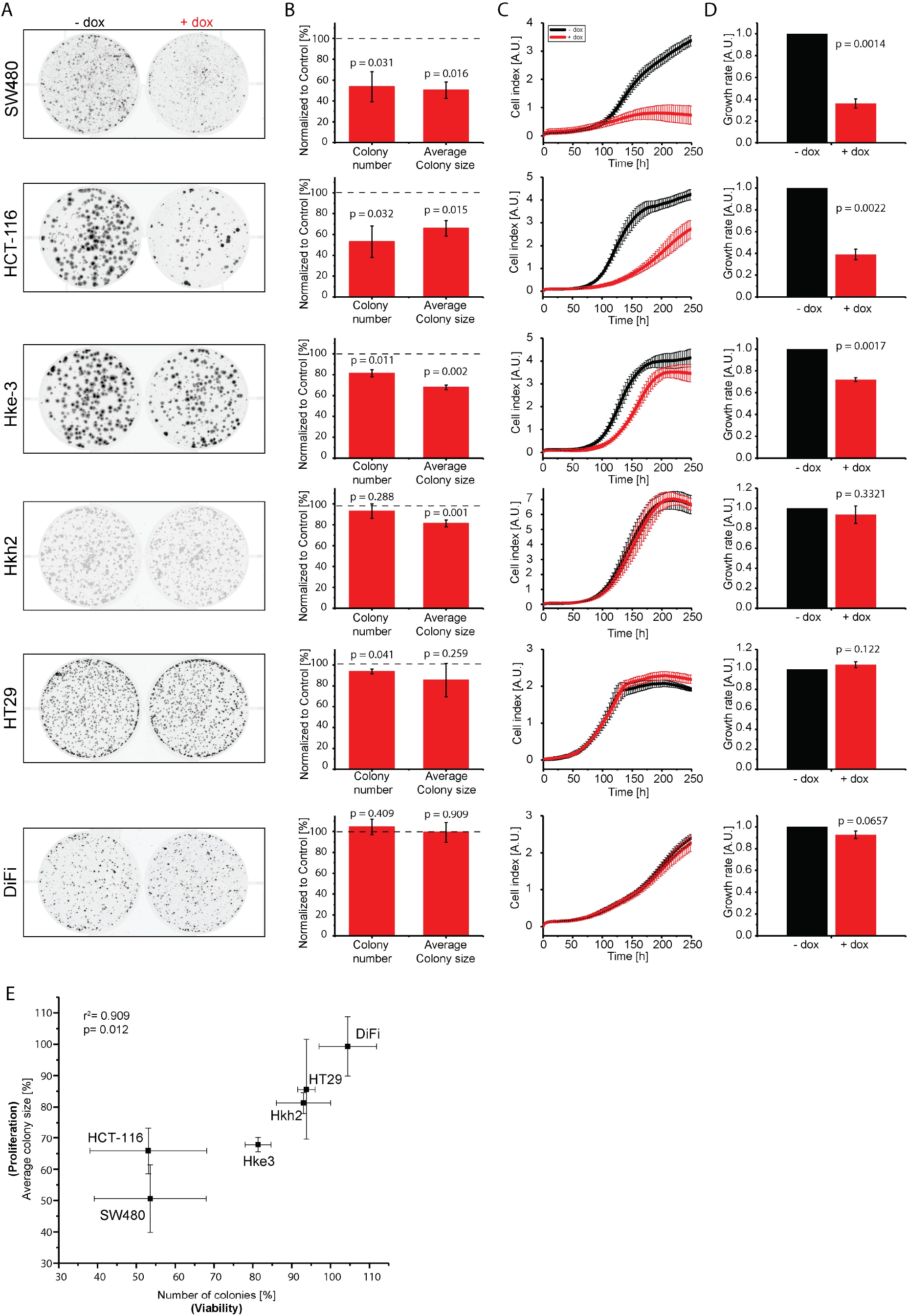
PDEδ knock down suppresses proliferation and survival of colorectal cancer cell lines harboring oncogenic KRas mutations. **(A)** Representative example out of three independent clonogenic assay experiments for the cell lines indicated. Cells were grown for ten days in the presence (+dox) or absence (-dox) of doxycycline. **(B)** Quantification of colony number ± s.d. and average colony size ± s.d. of three independent experiments. Knock down wells were normalized to the respective untreated control (dashed line). Significance was calculated using one sample *t* test. **(C)** Representative RTCA profiles out of three independent experiments. Cell indices ± s.d. of four replicates were measured in the presence (red) or absence (black) of doxycycline. Doxycycline was added at the beginning of the measurement. **(D)** Growth rates ± s.d in the presence (red) and absence (black) of doxycycline of three independent experiments. Growth rates were calculated by the area under curve over 240 h and normalized to the respective untreated condition. Significance was calculated using one sample *t* test. **(E)** Correlation plot of colony number ± s.d. versus average colony size ± s.d relative to respective control conditions under PDEδ knock down as determined in (B). Pearson’s correlation analysis shows a correlation of 0.909.

The clonogenic assays were complemented with real-time cell analysis (RTCA), where changes in the coverage of a surface by cells is measured by impedance (figure 2 C) [18]. Consistent with the clonogenic assays, PDEδ knock down resulted in a strongly reduced proliferation of CRC cell lines harboring oncogenic KRas, while the growth rates of KRas wild-type cell lines Hkh2, HT29 and DiFi were again comparable to the respective controls (figure 2 D). The CRC cells with heterozygote oncogenic KRas mutation (HCT-116 and Hke3) both exhibited reduced cell proliferation upon doxycycline induction. However, whereas Hke3 only exhibited a delay in cell proliferation after doxycycline administration, the rate of proliferation was affected in HCT-116, resulting in a substantially reduced cell number. In contrast, the growth rate of homozygote oncogenic KRas mutation bearing SW480 cells completely stagnated after doxycycline administration and cell death became apparent after 175 h from the negative growth rate (decrease in cell index).

We next compared the effects of two small-molecule PDEδ inhibitors with different chemotypes, Deltarasin [10] and Deltasonamide 2 [12] (figure 3 A, F), on growth rate and cell viability within the CRC cell panel. Both inhibitors competitively bind to the hydrophobic binding pocket of PDEδ as mediated by hydrogen bonds (H-bounds). However, while Deltarasin engages only in 3 H-bonds exhibiting a corresponding moderate affinity (K_D_= 38 ± 16 nM) [10], Deltasonamide 2 engages in 7 H-bonds and exhibits a high affinity (K_D_= 385 ± 52 pM) [12]. To determine the effects of the dose of inhibitors on proliferation, we performed RTCA measurements for cell growth (figure 3 B, G; supplementary figure 3) and flow cytometry based 7-AAD (7-Aminoactinomycin D) single cell fluorescence assays that report on cell death [27] (figure 3C,H; supplementary figure 4, 5). We related the effects of the inhibitors on proliferation and cell viability by determining EC_50_ values by sigmoidal curve fitting of the calculated growth rates against inhibitor dose and plotted these against the difference in cell viability at highest inhibitor dose in comparison to the DMSO control (Δ cell viability). In these inhibitor correlation plots (figure 3 D, I), SW480 exhibited the lowest EC_50_ (Deltarasin: 2.86 ± 0.31 μM, Deltasonamide2: 1.24 ± 0.06 μM) as well as highly compromised viability. The three isogenic cell lines (HCT-116, Hke3, Hkh2) showed comparable EC_50_ values, while cell viability of oncogenic KRas-lacking Hkh2 was less affected compared to HCT-116 and Hke3. The DiFi and HT29 cells that lack oncogenic KRas were clearly separated from oncogenic KRas harboring cell lines, where DiFi exhibited the highest EC_50_ (Deltarasin: 8.92 ± 0.7 μM, Deltasonamide2: 4.02 ± 1 μM) and HT29 viability was not affected by inhibitor administration. As expected, the high affinity inhibitor Deltasonamide 2 showed a shift to lower EC_50_ values for all CRC cell lines. Strikingly, the correlation plots of both Deltarasin (figure 3 D) and Deltasonamide 2 (figure 3 I) showed a similar alignment of cell lines with respect to their KRas mutation status and this alignment was reminiscent to that of PDEδ knock down (figure 2 E). This further strengthens the argument [10, 12] that the effect of the inhibitors on proliferation is due to specific targeting of PDEδ. To further compare dose-response profiles between Deltarasin and Deltasonamide 2, we plotted viability (7-AAD staining) versus growth rate (RTCA) in dependence of the respective inhibitor dose and cell line (figure 3 E, J). This again revealed the similarity in dose-response profiles between SW480, HCT-116 and Hke3 regarding reduced viability and growth for both inhibitors. The wild-type KRas cell lines were clearly less affected in both proliferation readouts, with the HCT-116-derived Hkh2 cells being the most responsive to the inhibitory effects on growth rate and viability.

**Figure 3.**
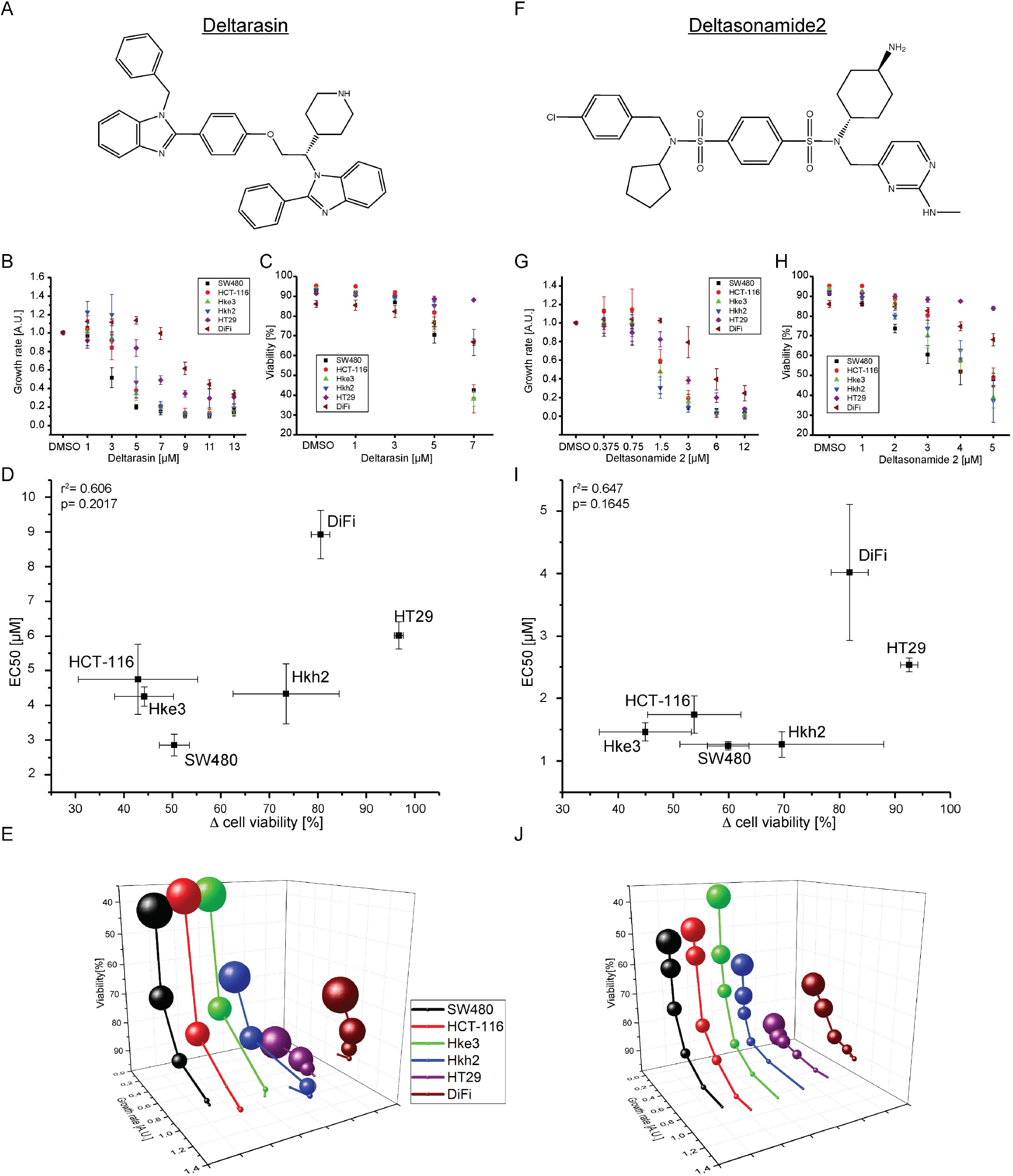
Dose-dependent inhibition of proliferation and viability reduction in human colorectal cancer cell lines by PDEδ inhibitors. **(A, F)** Chemical structures of the small molecule PDEδ inhibitors Deltarasin and Deltasonamide 2. **(B, G)** Growth rate ± s.d. in dependence of Deltarasin or Deltasonamide 2 dose. Growth rates were determined by integration of the area below the RTCA curves (Sup. Fig 3) over 60 h after drug administration and normalized to the DMSO control. **(C, H)** Cell viability ± s.d. in dependence of Deltarasin or Deltasonamide 2 dose in CRC cell lines after 24 h of drug administration. Cell death was determined by viability staining using 7-AAD. DMSO was used as vehicle control. **(D, I)** Correlation of Δ Cell viability ± s.d. versus EC_50_ ± s.d. for Deltarasin (D) and Deltasonamide 2 (I). Δ cell viability was calculated between DMSO control and the highest used inhibitor concentration, respectively. EC_50_ values were determined by sigmoidal curve fit of the growth rates depicted in B and G. **(E, J)** Four-dimensional correlation of growth rate and cell viability in dependence of inhibitor dose and CRC cell line for Deltarasin (E) and Deltasonamide 2 (J). The dot size is proportional to the applied inhibitor concentration.

Both PDEδ knock down and small-molecule inhibition were most effective in SW480 cells, which are homozygote for oncogenic KRas [19]. SW480 also exhibited the highest expression levels of Ras and PDEδ. Together, this implies that proliferation and survival of SW480 are depending on oncogenic KRas (and thereby PDEδ) and that these cells have no compensatory mechanism to rescue for PDEδ loss or inhibition. The isogenic cell lines HCT-116, Hke3 and Hkh2 are a well-suited system to study effects of PDEδ interference in a presumably isogenic background since they should only differ in their KRas mutation status [20]. Our results (figure 1 C) however showed that Hke3 cells still possess GTP-loaded Ras under serum-starved conditions and thereby confirmed that they still harbor an oncogenic KRas mutation [21]. In contrast, the oncogenic allele was successfully removed in the Hkh2 cell line, manifested in the low level of detected Ras-GTP (figure 1 C). Indeed, the parental HCT-116 cell line showed a stronger reduction in cell growth and viability by PDEδ knock down or inhibitor treatment compared to Hkh2. In contrast, effects on growth rate and cell viability were comparable between HCT-116 and Hke3 after PDEδ inhibition, whereas cell viability of Hke3 was less affected by PDEδ knock down. This indicates that the oncogenic KRas expression levels are an important determinant for cell survival but less for proliferation in CRC cells. This would also point at that oncogene addiction is related to the expression level of the oncogene. In this context, the reported correlation between increased Ras expression levels and oncogenic KRas mutations [28] was also apparent within our CRC cell panel.

Strikingly, the proliferation and survival of BRaf(V600E) bearing HT29 [17] and EGFR overexpressing DiFi [23, 24] was not affected by PDEδ knockout and were the least sensitive to both tested PDEδ inhibitors. In the MAP kinase signaling network [29], BRaf is activated downstream of KRas, making those cells that harbor the BRaf(V600E) mutation independent of KRas signal input and thereby its localization. This is consistent with PDEδ interference not affecting the proliferation of these cells. However, DiFi cells feature an up-regulated EGFR expression level [23, 24] and EGFR is located upstream of KRas in the MAP kinase signaling network. One would therefore assume that PDEδ down modulation would affect signal propagation in the MAPK network in these cells, which was not the case. However, DiFi cells also exhibit low levels of Ras protein (figure 1B), and it is therefore likely that other signals emanate from overexpressed EGFR, possibly via the PI3K-Akt axis, that sustain proliferation and survival. Both DiFi and HT29 cells, expressed the lowest amount of PDEδ as well as Ras proteins among our tested CRC cell lines, and PDEδ expression level was correlated to oncogenic Ras activity. This indicates the interdependence of oncogenic Ras activity and the solubilizing activity of PDEδ that can be exploited to affect oncogenic KRas signaling in cancer cells by inhibition of PDEδ. Indeed, small molecule inhibition of PDEδ in these CRC cell lines phenocopied PDEδ knock down. The latest generation of high affinity PDEδ inhibitors such as Deltasonamide 2 thereby proved to be the superior inhibitor [12]. The discrepancy between the μM concentration of Deltasonamide 2 that induce a growth inhibitory effect and its K_D_ for PDEδ (~385 pM) is due to its low partitioning in the cytosol. However, our results show that potent inhibitors of the KRas-PDE6δ interaction might impair the growth of CRC driven by oncogenic KRas and may offer new therapeutic angles for colorectal cancers harboring oncogenic KRas mutations that are unresponsive to treatment [14, 15].

## Acknowledgements

This research was supported by the Deutsche Krebshilfe (Grant 110995) and the European Research Council (ERC Grant 322637).

## Author contributions

P.I.H.B. conceived the project. D.C.T. and C.H.K. generated stable inducible shRNA-PDEδ cell lines. C.H.K. and D.C.T. performed western blot analysis. C.H.K. performed clonogenic assays and J.H. and C.H.K. analyzed the data. C.H.K. and H.A.V. performed real-time cell analysis measurements. C.H.K. performed viability assays. S.M. and P.M.G. synthesized Deltasonamide 2. C.H.K. and P.I.H.B. wrote the manuscript.

---

**Supplementary figure 1:**
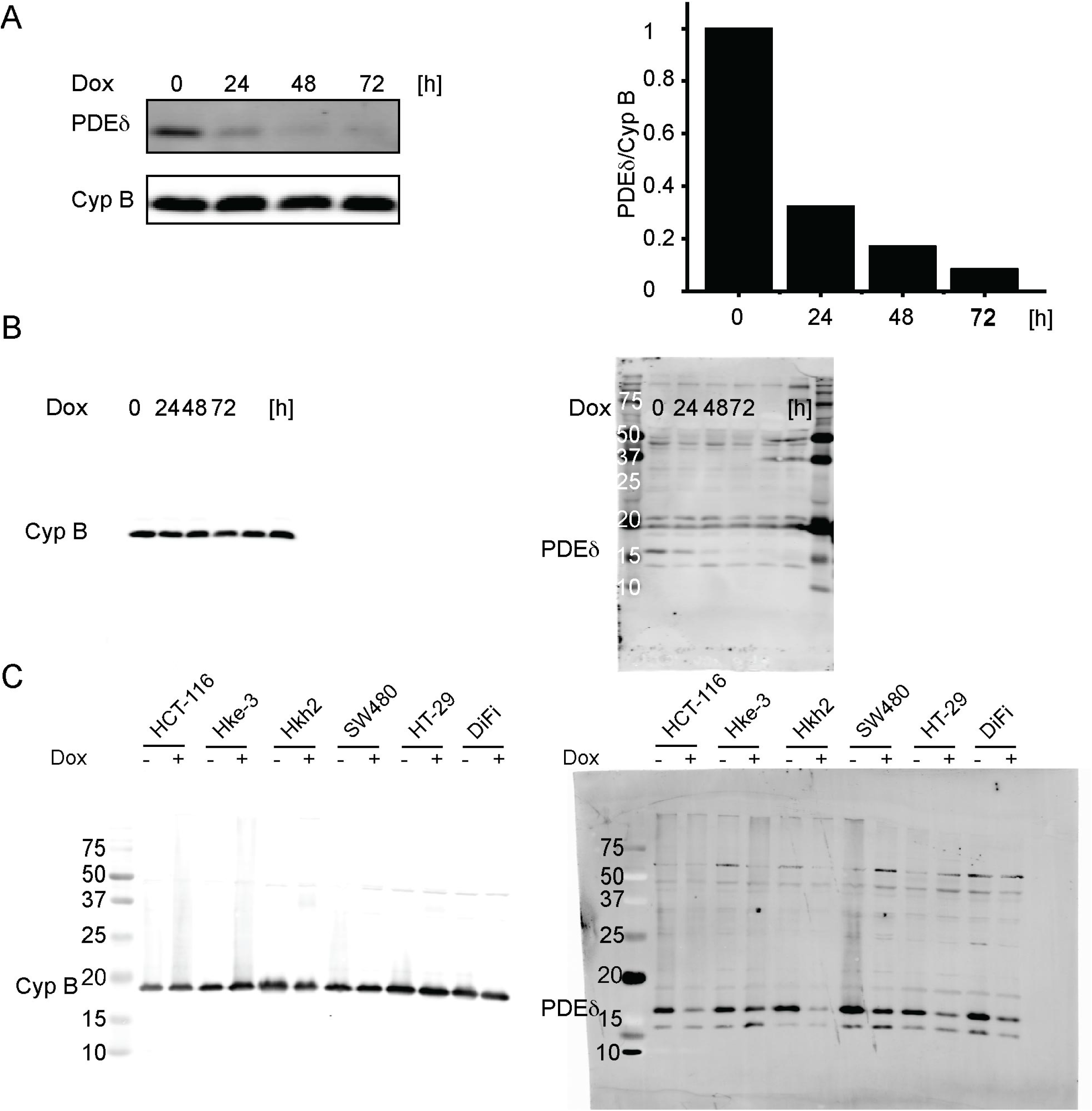
**(A)** Left: PDEδ protein level in Hke3 cells after increasing doxycycline administration periods determined by western blot. Cyclophilin B was used as loading control. Right bar graph: quantification of PDEδ protein levels normalized to the untreated control. **(B)** Uncropped western blot used for (A). **(C)** Uncropped western blot used for inset and quantification in figure 1 A.

**Supplementary figure 2:**
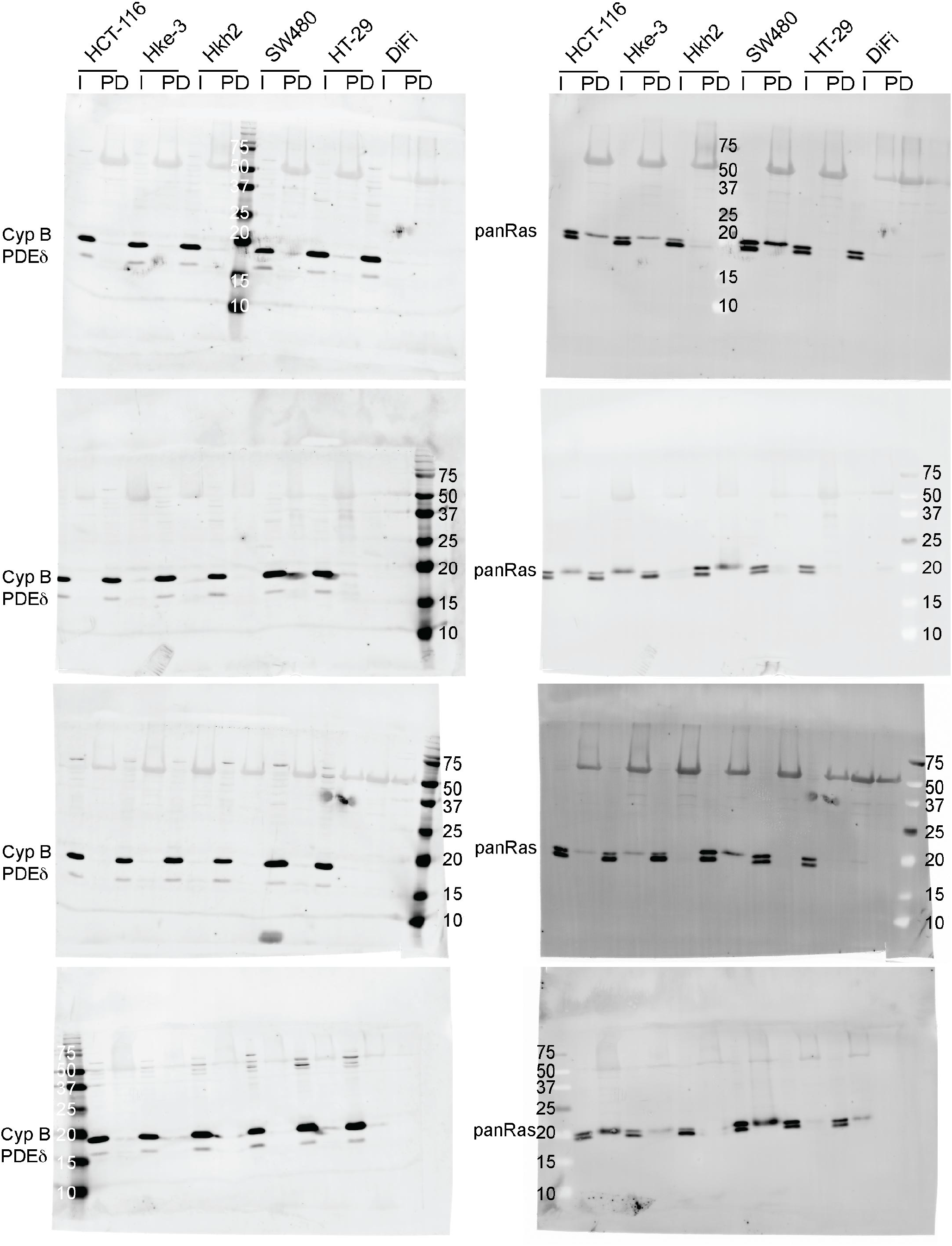
Uncropped western blots (n=4) used for inset and quantitative analysis in figure 1 B.

**Supplementary figure 3:**
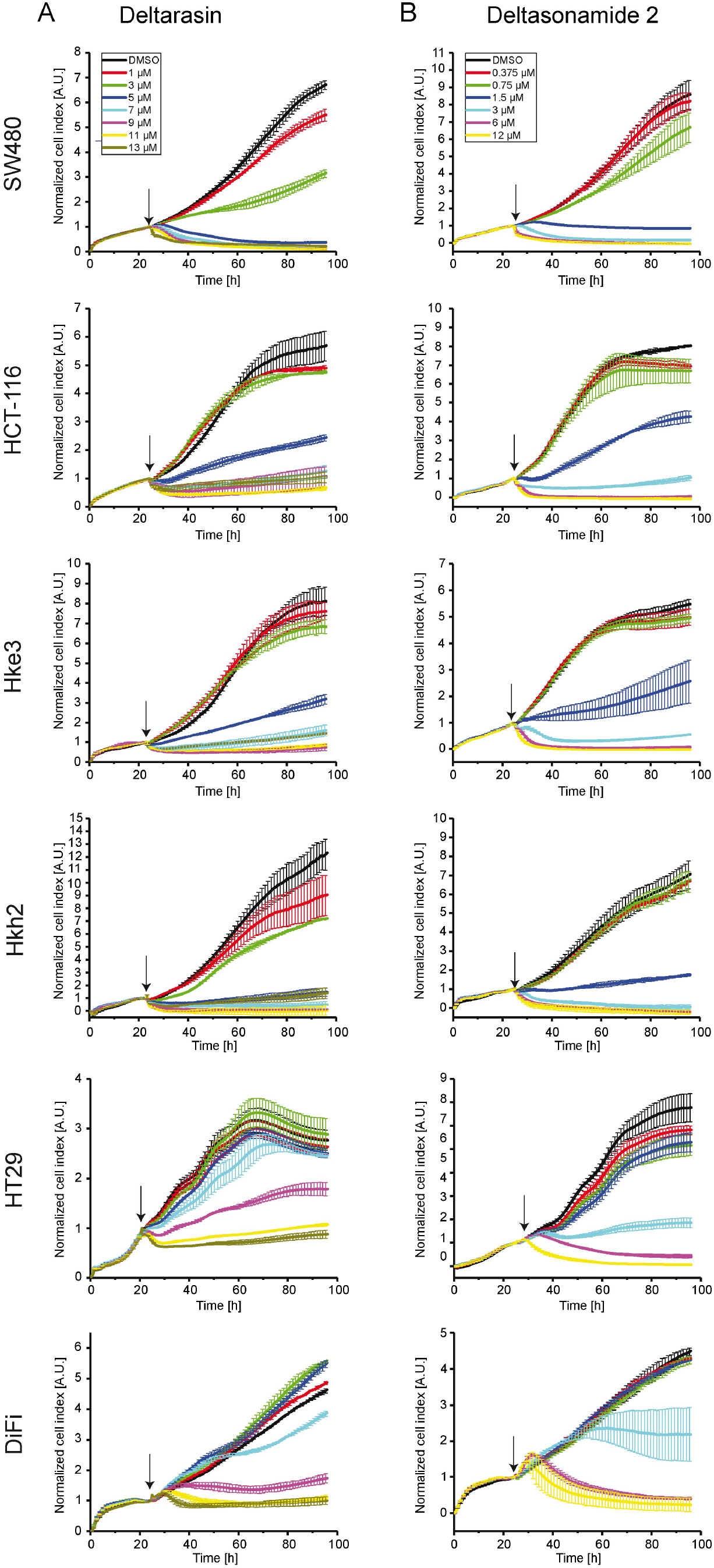
**(A) – (B)** Representative RTCA profiles (n=3) of colorectal cancer cell lines with distinct KRas mutation status treated with different doses of Deltarasin (A) or Deltasonamide 2 (B). Cell indices ± s.d. were measured in duplicates and normalized to the time point of drug administration (arrow).

**Supplementary figure 4:**
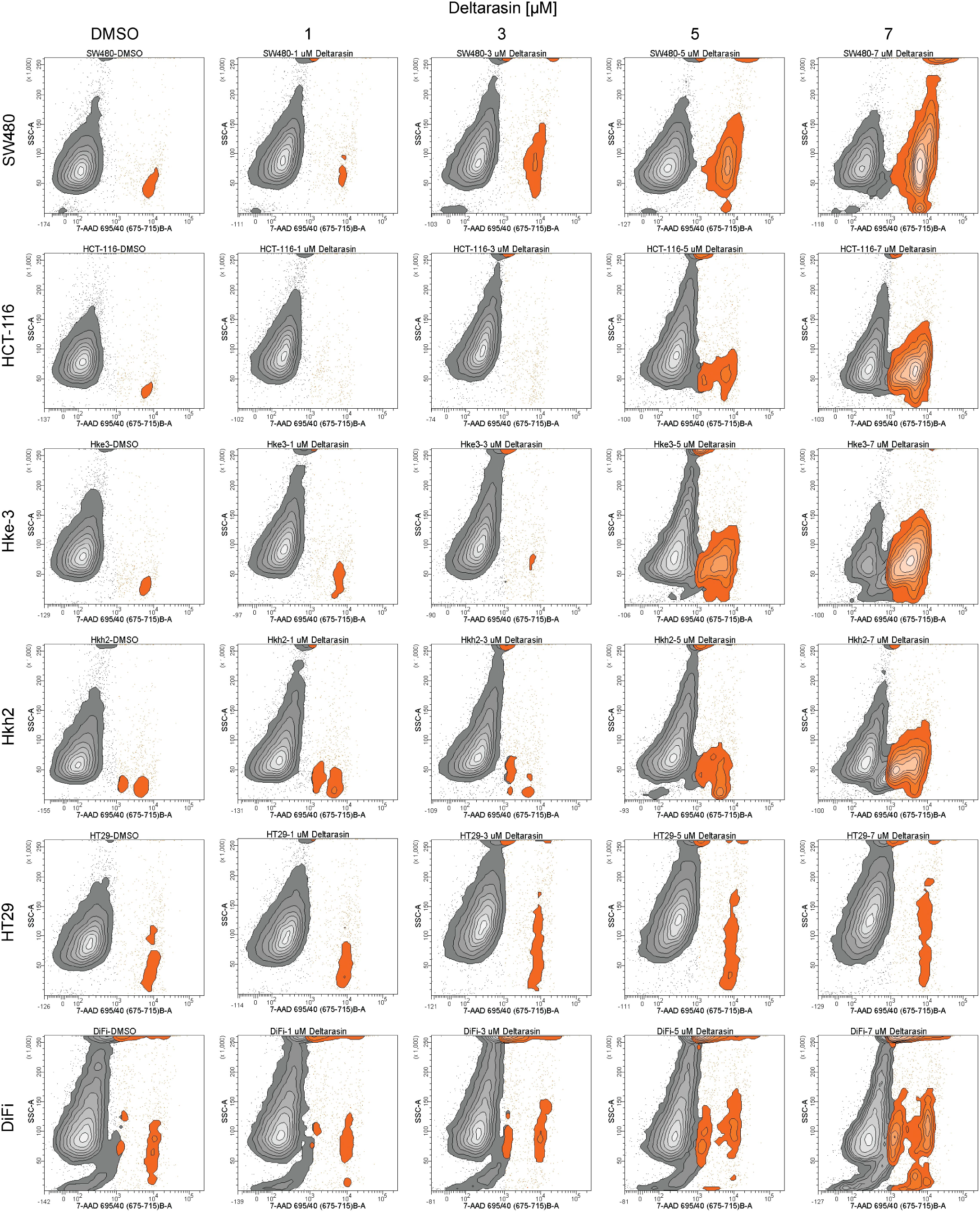
Representative contours plots (n=3) of side scattering (SSC) versus 7-AAD fluorescence of CRC cells treated with different doses of Deltarasin. 7-AAD negative cells are shown in black, 7-AAD positive cells are shown in orange. Viable, 7-AAD negative cells (black) were gated based on unstained control cells.

**Supplementary figure 5:**
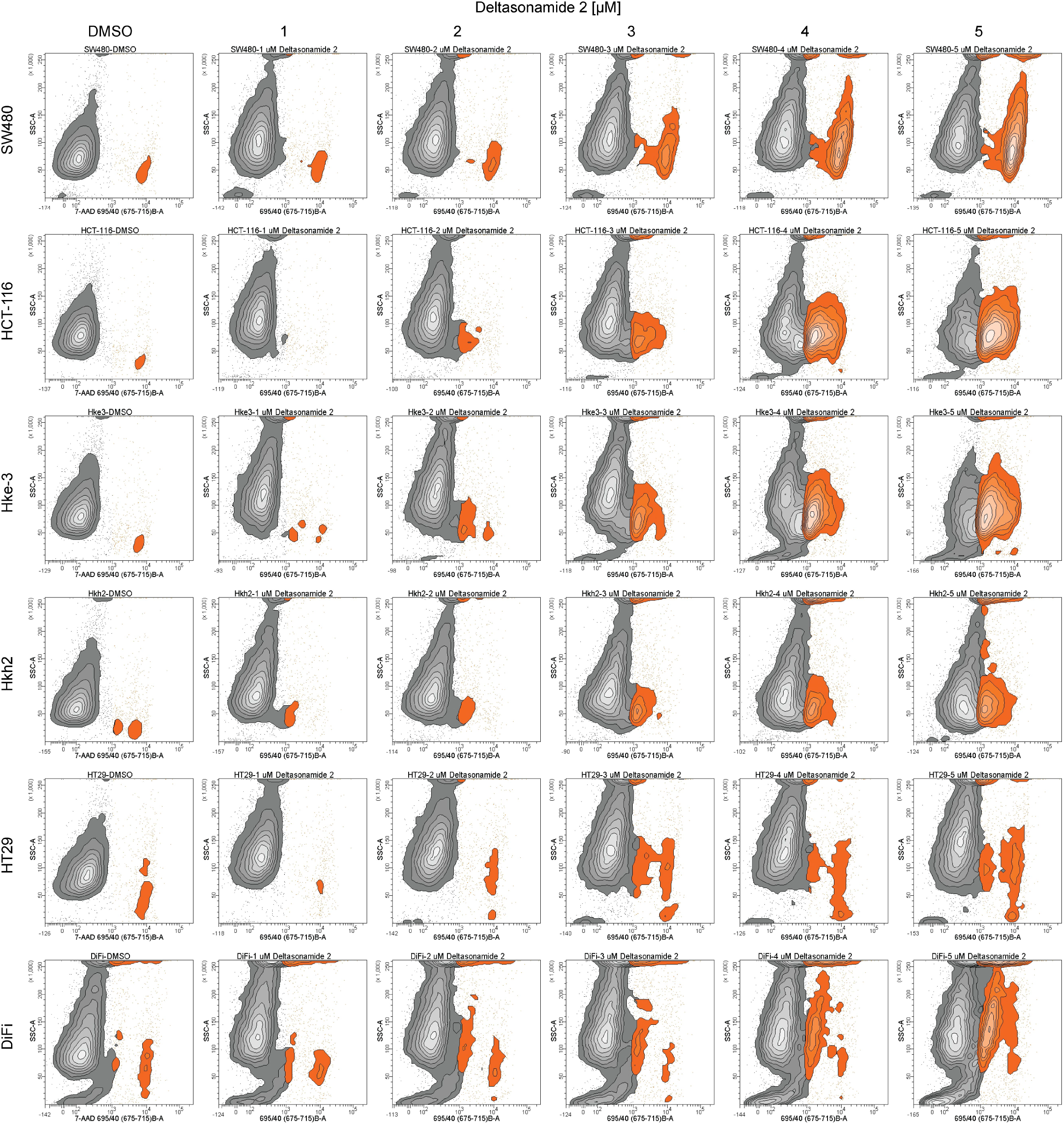
Representative contours plots (n=3) of side scattering (SSC) versus 7-AAD fluorescence of CRC cells treated with different doses of Deltasonamide 2. 7-AAD negative cells are shown in black, 7-AAD positive cells are shown in orange. Viable, 7-AAD negative cells (black) were gated based on unstained control cells.

## Abbreviations

CRC: colorectal cancer
PM: plasma membrane
GDI: guanine nucleotide dissociation inhibitor
RE: recycling endosome
hPDACs: human pancreatic ductal adenocarcinoma cells
shRNA: short hairpin RNA
RTCA: real-time cell analysis
7-AAD: 7-Aminoactinomycin D

